# Colonization of 3D-organotypic human skin by the Lyme Disease pathogen, *Borrelia burgdorferi*

**DOI:** 10.64898/2026.07.20.739593

**Authors:** Jaxon J. Kramer, Amanda J. Brinkworth

## Abstract

*Borrelia burgdorferi* is a bacterial pathogen transmitted by ticks that is the causative agent of Lyme Disease. When studying interactions between *B. burgdorferi* and skin following a tick-bite, existing models consist primarily of murine skin which has different cellularity and thickness than human skin or *ex vivo* human biopsies that can be difficult to obtain and can have high variability. This presents a need for a reproducible human skin model. Herein, we adapt an existing human organotypic skin model that is simply composed of dermal fibroblasts and stratified epidermal keratinocytes and develop an infection assay mimicking skin reinfection to characterize *B. burgdorferi* colonization. Normal spirochete morphology and a stressed *B. burgdorferi* morphology known as a “round body” were observed. Peak *B. burgdorferi* invasion was observed at 24 hours (h) with peak round body formation at 48 h. By breaking the skin down into its individual components, we observed an increase in the number of round bodies in the presence of dermal fibroblasts. The presence of keratinocytes or extracellular matrix alone had no effect on round body formation, indicating a dermal fibroblast-mediated mechanism. We also demonstrate tissue-to-tissue dissemination and colonization, setting the groundwork for future studies with other tissues. Collectively, these results prove that this is a valid human skin model to study *B. burgdorferi* colonization during secondary dissemination.

## Introduction

*Borrelia burgdorferi* is a spirochete and causative agent of Lyme disease (LD). LD is one of the most prevalent tick-borne diseases in the world and in the United States, with nearly an estimated half million cases annually and increasing each year [1,2]. The economic burden of LD in the United States has ballooned with an estimated $786 million spent annually on diagnosis and treatment [3]. LD manifestations primarily include the presence of the *erythema migrans* lesion, otherwise known as the “bullseye rash” days to weeks following a tick-bite and *B. burgdorferi* transmission [4–6]. If left untreated, chronic, secondary forms of LD can arise following dissemination to distant tissues, including multiple *erythema migrans*, carditis, arthritis, and neuroborreliosis [4,5].

*B. burgdorferi* is transmitted via *Ixodes* tick species and maintained in an enzootic cycle primarily with *Peromyscus* mouse species, which can be a reservoir for *B. burgdorferi* [7]. Within the enzootic cycle, all three life stages (larval, nymphal, and adult) can transmit and acquire *B. burgdorferi*, but nymphs account for the majority of transmission events to humans [2,7]. Upon transmission into human skin, *B. burgdorferi* will encounter a variety of host immune factors and cells. These include keratinocytes, fibroblasts, endothelial cells, CD4^+^ and CD8^+^ T cells, γδ T cells, B cells, mast cells, macrophages, dendritic cells, and Langerhans cells [8–10]. Upon inflammation, additional immune cells including monocytes, granulocytes, and additional lymphocytes are recruited to clear debris, prevent infections, and facilitate the wound healing process [9]. During tick feeding, *B. burgdorferi* enters skin alongside tick saliva, which contains a cocktail of molecules that modulate the host immune response to the benefit of both the tick and pathogen [11,12].

Current models to study the vector-*B. burgdorferi*-skin and *B. burgdorferi*-skin interfaces primarily use murine models. Among the different lab strains, C3H/HeN mice have been identified as more sensitive to infection in comparison to BALB/c and C57BL/6 mice and exhibit a greater pro-inflammatory response [13,14]. Other animal models used include New Zealand White rabbits [15] and rhesus monkeys [16]. While rhesus monkeys presented with *erythema migrans* and associated chronic manifestations of LD such as Lyme arthritis and neuroborreliosis [16], less studies have been performed due to animal size, care, cost, and ethical issues [7]. Outside of animal models, *ex vivo* human skin has been used as well to study vector-pathogen-host interactions [17–19], but skin availability and patient variability issues remain a barrier to many studies. Biopsies from human *erythema migrans* have also been analyzed for the immunological markers and cells that are recruited in the days to weeks following tick transmission of *B. burgdorferi* [20]. A recent study analyzed early infection in events from recent tick-bite, revealing a key role of different CD4^+^ and CD8^+^ T cell subsets in skin responses by single cell RNA-sequencing [20]. However, the complexity of human skin, the small proportion of immune cells compared to fibroblasts and keratinocytes [20], and potential for crosstalk between multiple cell types makes interpreting these results challenging.This presents a need for a reproducible human skin model that can be additive, while still maintaining skin architecture. One option is a 3D-organotypic human skin model, which has existed for over 40 years [21,22]. The simplest version of a complete organotypic skin model includes a dermal layer that consists of extracellular matrix (ECM) and fibroblasts along with an epidermal keratinocyte layer on top that is stratified at an air-liquid interface (ALI). Organotypic human skin has been used for other pathogen studies [23–25] and represents a standardized, reproducible model that is scalable and adaptable. Importantly, in the organotypic model, additional cell types or ECM components can be added or subtracted to dissect specific phenotypes [26–28].

Herein, we have adapted an existing organotypic human skin model [29] with a goal to study interactions between *B. burgdorferi* and resident skin cells. We have developed a secondary dissemination assay (SDA) that mimics skin reinfection by allowing for spirochete migration from the dermal side. Both normal spirochete-shaped and stressed round body forms (RBs) [30–32] were observed. Peak levels of *B. burgdorferi* were observed in the skin at 24 h post-inoculation while the highest proportion of *Borrelia* in stressed RB form was observed at 48 h and beyond. *B. burgdorferi* were more abundant in the dermal compartment compared to the epidermis, with RB formation being similar in the two compartments.

By systematic dissection of the organotypic skin components, we determined that RB formation depends on the presence of dermal fibroblasts, while the presence of epidermal keratinocytes and varying ECM content had no effect. Dermal fibroblasts alone also had a negative effect on total numbers of bacteria, which was relieved by the presence of keratinocytes. We also prove that RB formation was not due to excess ROS in our model. Overall, these data validate an organotypic human skin model for *B. burgdorferi* infection that supports growth, dissemination, and persistent forms, making it highly relevant for future host-pathogen interaction studies.

## Results

### Establishment of human organotypic skin for *B. burgdorferi* infection

Current models to study primary *B. burgdorferi* infections in the skin include both murine [33–35] and human [17–19] skin. Murine and human skin have quite a few differences, including thickness, epidermal layers, number of hair follicles, and immune cell populations, among others [36,37]. All of these differences can play a role in the response to a tick-bite and pathogen transmission. *Ex vivo* human skin is the gold standard, but accessibility and reproducibility can prove difficult. This presents a need for a relevant and replicable human skin model to study *B. burgdorferi* colonization and interaction with epidermal and dermal resident cells.

Our model has been adapted from a previous 3D-organotypic skin model [29]. Briefly, dermal fibroblasts are embedded into a 4 mg/mL type I collagen layer. After gel solidification at 37□, HaCaT keratinocytes are seeded on top and left submerged for 2 days to reach confluency. Afterwards, media is removed from the top to create an ALI and induce stratification. For 10-15 total days, media is changed every 2 days in the lower chamber to refresh necessary growth factors.

We monitored the maturation of organotypic skin in OCT-embedded cross-sections by immunofluorescence. On Day 10, epidermal and dermal layers are easily identifiable via wheat germ agglutinin (WGA) staining (Fig. 1A). Ki67 staining marks cells actively proliferating cells, particularly at the basal epidermal layer and the dermal fibroblasts at both Days 10 and 15 (Fig. 1A). Similarly, Cytokeratin 14 (CK14) staining marks proliferating keratinocytes and is apparent in both Day 10 and 15 skin (Fig. 1A). Along with this, loricrin staining in the *stratum corneum*, the outermost layer of the epidermis, is more defined and apparent on Day 15 compared to Day 10 (Fig. 1B). Similarly, filaggrin deposition started by Day 15, which is a key component of the epidermal surface barrier.

**Figure 1.**
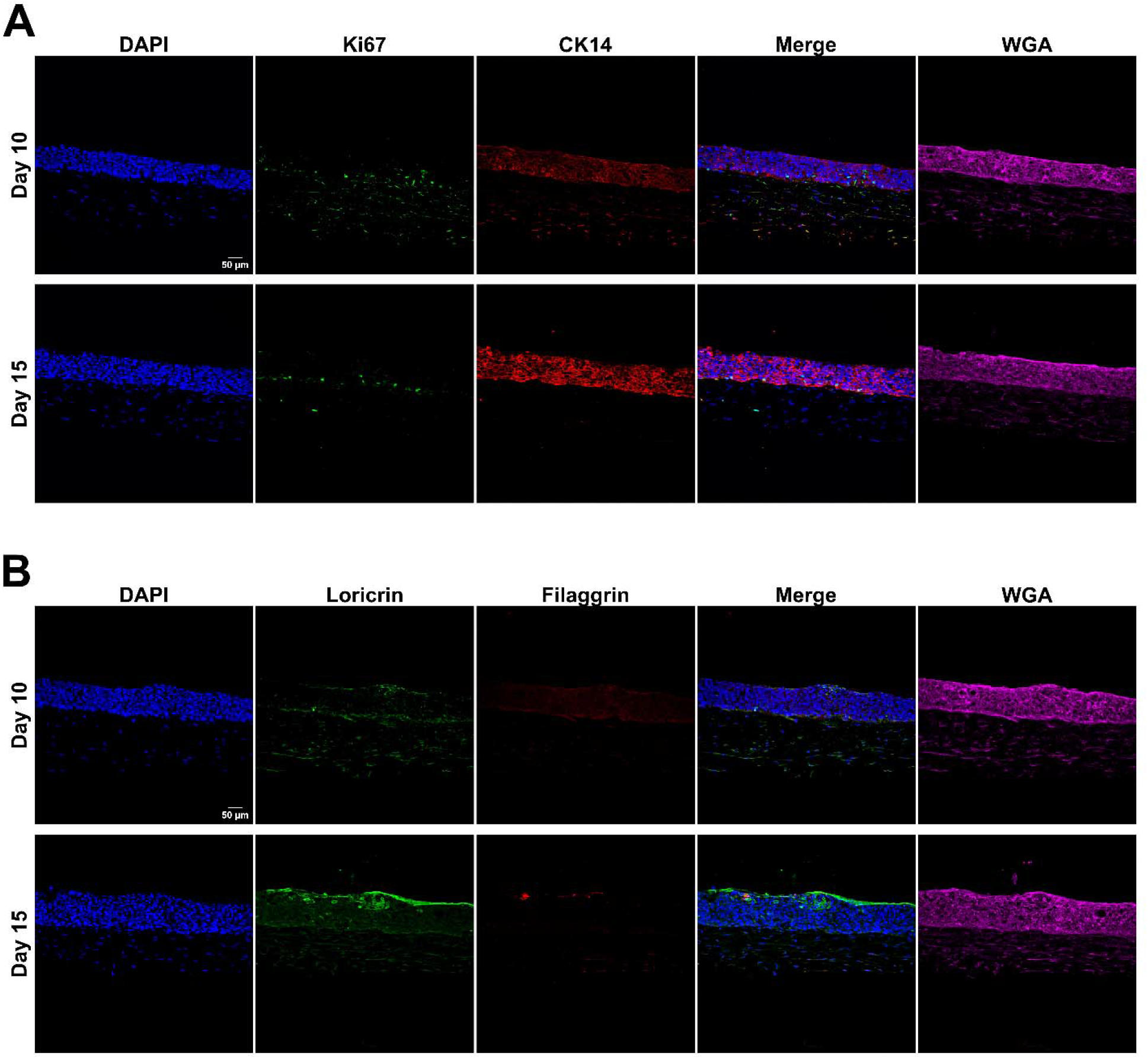
Day 15 organotypic skin represent a fully differentiated human skin model. (A) Representative images of Day 10 and 15 organotypic skin are stained with DAPI (blue) to mark the nucleus, Ki67 (green) to mark proliferating cells, particularly in the basal epidermal layer, CK14 (red) to mark cytokeratin 14, and wheat germ agglutinin (WGA) to differentiate all cellular layers. (B) Representative images of Day 10 and 15 organotypic skin are stained with DAPI (blue) to mark the nucleus, loricrin (green) to mark the stratum corneum, filaggrin (red) to mark the stratum corneum, and WGA to differentiate skin layers.

Taken together, Day 10 organotypic skin can be used to study the effects of a co-culture model in response to infection while Day 15 organotypic skin represents a tool to study a primary infection due to a completely differentiated and stratified epidermis.

### Secondary dissemination assay presents a new way to colonize tissue

After initial skin infection in mice, *B. burgdorferi* can disseminate through the blood to other tissues and organs [38–42]. This includes reaching skin and distal sites such as joints, heart, liver, and spleen [38,40,42]. Most, if not all, studies use mice for secondary dissemination studies, necessitating a reproducible human skin model to monitor colonization. To determine optimal infection conditions, we generated a medium that is beneficial to the skin as well as *B. burgdorferi*. To initially test these growth conditions, 3×10^6^ spirochetes were inoculated into BSK-H alone, 1:1 BSK-H and Fibroblast Media (FBM), or FBM alone. Early log phase growth was achieved by Day 4 in BSK-H while 1:1 BSK-H and FBM grown spirochetes began increasing starting at Day 5 (Fig. S1A). Stationary phase was reached by Day 7 in the BSK-H culture while it was reached on Day 6 for the 1:1 medium (Fig. S1A). Across all time points, the FBM only culture never supported spirochete growth (Fig. S1A). Using motility as a proxy for spirochete viability, percent motility was calculated with over 90% of spirochetes being motile until Day 4 in BSK-H, before gradually decreasing as the culture reaches stationary phase (Fig. S1B). Across all time points, at least 50% of spirochetes were motile in the 1:1 medium while the FBM only culture had little to no motility after Day 4 (Fig. S1B). Taken together, this indicates that the 1:1 media can support *B. burgdorferi* growth and health for organotypic skin inoculation studies, albeit with slower growth and reaching stationary phase at a lower density than BSK-H media alone.

Using the organotypic skin model, we place 2 x 10^6^ *B. burgdorferi* in the 1:1 media beneath the organotypic skin (Fig. 2A) and leave it static to enable dissemination. Since the skin requires a media change once every 48 h, 24 h was chosen as the time point for the initial media change (Fig. 2B). An initial time course revealed that there was an increase in *B. burgdorferi* invasion up to 24 h before a media change occurred (Fig. 2C, D, and F). At each time point, both *B. burgdorferi* spirochetes and RBs, an alternative stressed morphology, were observed in skin cross-sections (Fig. 2C). This morphology has been previously associated with osmotic shock, nutrient deprivation, presence of specific antibiotics, and oxidative stress, among others [43–45]. RBs have also been previously identified in *in vivo* skin biopsies from Lyme Disease patients [46,47]. In our assay, peak RB formation was observed at 48 h (49.7%), 24 h after the media change (Fig. 2C and E). Spirochetes were also quantified from the media below the skin (Fig. 2G). As expected, the average number of *B. burgdorferi*/field in the media below dropped after the 24 h media change, though spirochetes were seen at both the 48 and 72 h time points, indicating continued movement of viable spirochetes in and out of the skin (Fig. 2G). At 24 h and beyond, more *B. burgdorferi* on average were observed in the dermis compared to the epidermis (Fig. 2H). RB formation was similar at each of these time points as well, with the percent of RBs appearing in the dermis being 29.3%, 51.7%, and 45% for 24, 48, and 72 h, respectively (Fig. 2I). This is compared to 26.7%, 38%, and 32.7% RB formation in the epidermis at 24, 48 and 72 h, respectively (Fig. 2I).

**Figure 2.**
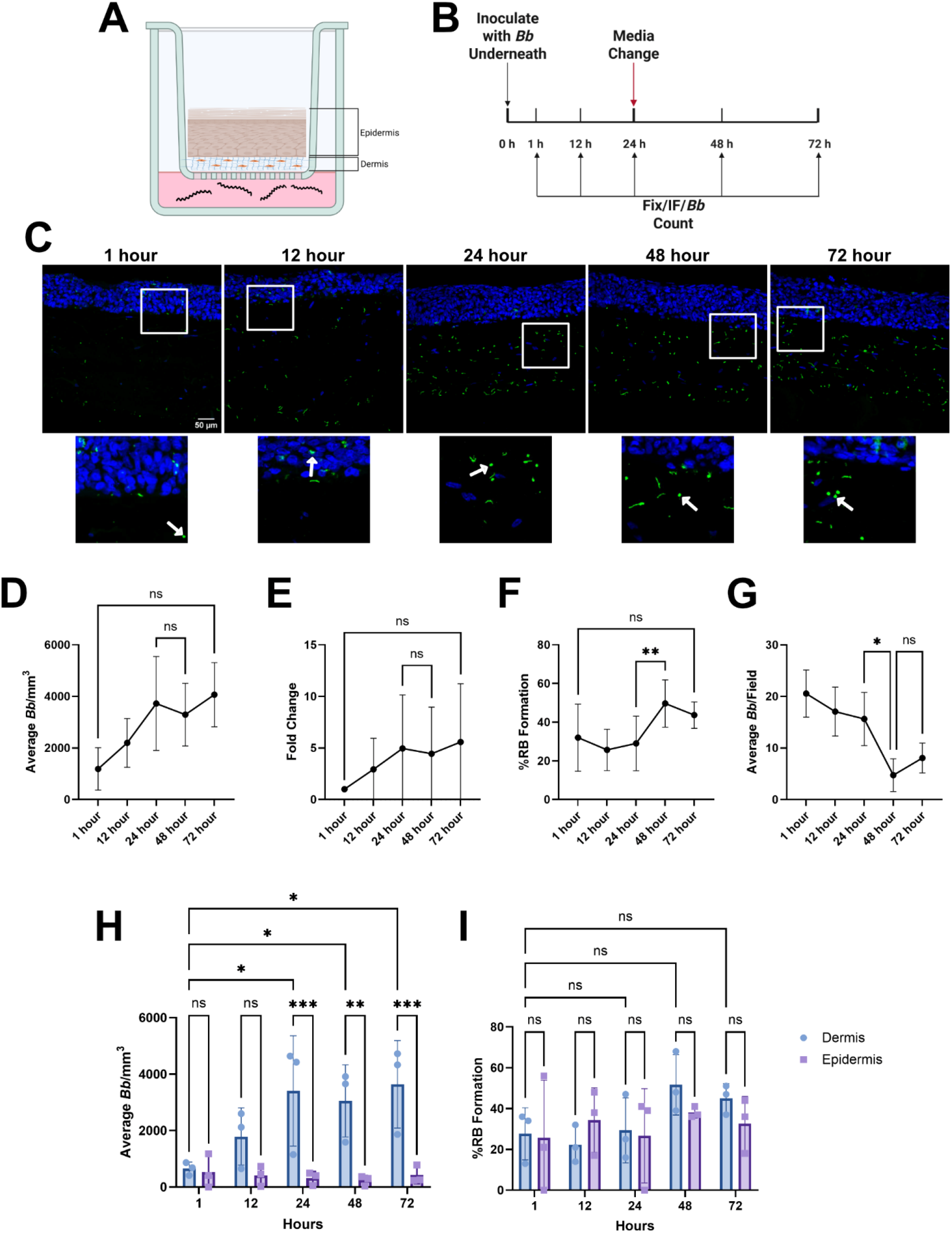
Round bodies (RBs) are identifiable across time course using a secondary dissemination assay. (A) *B. burgdorferi* is placed in a 1:1 mix of BSK-H and fibroblast media below the organotypic skin and left static for up to 72 h. (B) Timeline of infection. At each indicated time point, infected skin was fixed, stained for *B. burgdorferi*, and counted. Media is changed at 24 h to sustain skin viability. (C) *B. burgdorferi* spirochetes (green, anti-*B. burgdorferi* FITC-polyclonal) and round bodies (RBs) are identifiable in 30-micron cross-sections by confocal microscopy. White arrows point to RBs at each time point. (D) After quantification, *B. burgdorferi* is normalized to a mm^3^ for total *Borrelia*. (E) Fold change of total *Borrelia* compared to 1 h in (D). (F) Percent round body formation (%RB formation) is calculated by counting the number of round bodies in the image and dividing by the total number of *Borrelia*. (G) Spirochetes were counted in the media below after each time point and expressed as Average *B. burgdorferi* (*Bb*)/Field. (H) Total *Borrelia* are calculated for the dermis and epidermis. (I) %RB formation is calculated for the dermis and epidermis. Each dot represents the average of a biological replicate. Experiments were performed in biological triplicate and technical duplicates. One-way ANOVA with Šídák’s multiple comparisons test was used to determine statistical significance between selected time points in (D), (E), (F), and (G) with GraphPad prism. Two-way ANOVA with Tukey’s multiple comparisons test was used to determine statistical significance between time points and spirochete location in (H) and (I). Error bars represent standard deviation. Significance is denoted as \**p* < 0.05 and \*\**p* < 0.01.

An extended time course out to 144 h revealed that *B. burgdorferi* is present in the organotypic skin long-term (Fig. S2B, C, and D). At these later time points, high amounts of RBs are observed with 55% and 54% being RBs at 96 and 144 h, respectively (Fig S2E). Other stressed morphologies such as “blebs” and degraded forms were also identified at 96 and 144 h (Fig. S2B). Similarly to earlier time points, majority of *B. burgdorferi* exist in the dermis at 96 and 144 h (Fig. S2F) while RB formation is significantly increased at 144 h compared to 1 h (55% vs. 9.5%) (Fig. S2G).

### ECM components do not alter *B. burgdorferi* colonization in organotypic skin

It is known that *B. burgdorferi* expresses a variety of adhesins that specialize in binding specific ECM components such as decorin, laminin, dermatan sulfate, and fibronectin and play a pivotal role in virulence [48–53]. MaxGel (Sigma), an *in vitro* human-derived basement membrane extract, was included in the dermis to further mimic the ECM content by adding additional collagen species, fibronectin, laminin, as well as various proteoglycans and glycosaminoglycans to potentially increase bacterial invasion. In each condition, addition of MaxGel had little to no effect on the total *B. burgdorferi* in the skin (Fig. S3A and B), as well as having no apparent effect on RB formation at 24 h (Fig. S3C).

### Secondary dissemination assay reveals dermal fibroblast-mediated stress

By breaking down the skin into its individual components, we can determine the source of stress leading to RB formation. In the Collagen Only (no cells) condition, we observed an increase in *B. burgdorferi* invasion (1.32-fold increase) (Fig. 3A and C) and decreased RB formation (17.3% vs. 37%) (Fig. 3D) compared to the complete skin condition. In the Collagen + HaCaTs (epidermis only) condition we observe comparable amounts of *B. burgdorferi* invasion (Fig. 3A, B, and C) and RB formation at 24 h (Fig. 3D). In the Collagen + NHDFs (fibroblasts only) condition, we observed significantly decreased *B. burgdorferi* invasion (2.82-fold decrease) (Fig. 3C) while observing comparable RB formation (36.7% vs. 37%) (Fig. 3D). This implies an NHDF-mediated RB formation mechanism.

**Figure 3.**
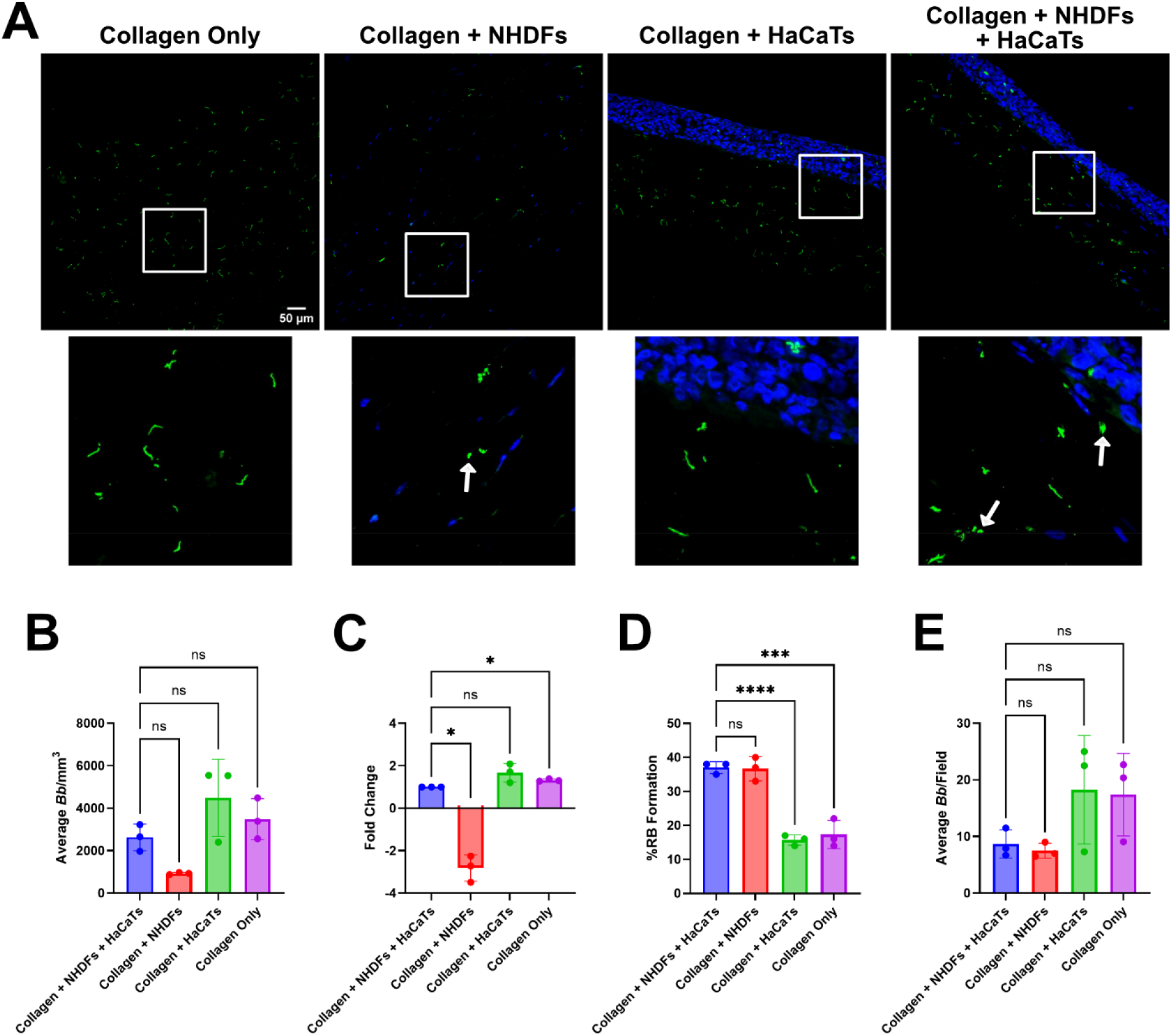
Dermal fibroblasts contribute to round body formation. (A) Representative images of organotypic skin without cells (Collagen Only), fibroblasts only (Collagen + NHDFs), keratinocytes only (Collagen + HaCaTs) and a complete model (Collagen + NHDFs + HaCaTs) that were infected using a secondary dissemination assay. White arrows point to RBs. (B) Total *B. burgdorferi* were quantified at 24 h in each skin condition. (C) Fold change of total *Borrelia* compared to the complete model in (B). (D) % RB formation was quantified at 24 h in each skin condition. (E) Spirochetes were counted in the media below in each condition and expressed as Average *Bb*/Field. Experiments were performed in biological triplicates and technical duplicates. Each dot represents the average of a biological replicate. One-way ANOVA with Dunnett’s multiple comparisons test was used to compare each condition to our control condition (Collagen + NHDFs + HaCaTs). Error bars represent standard deviation. Significance is denoted as \**p* < 0.05, \*\*\**p* < 0.001 and \*\*\*\**p* < 0.0001.

Dermal fibroblasts have been shown to play a role in the immune response to *B. burgdorferi,* including upregulation of cytokines and chemokines at the transcript (transforming growth factor-beta, platelet-derived growth factor-alpha, CCL2, CXCL10, interleukin-6 (IL-6), IL-8, type I interferons) [54–56] and protein (IL-6) [55] levels. It is worth noting that the Collagen + NHDFs condition presents as fibrotic-like and contracted compared to the complete skin condition (Fig. S4A), which may serve as a physical barrier preventing effective spirochete migration into the skin (Fig. 3A). *B. burgdorferi* was counted in the media below in each condition after 24 h and a decrease in the average spirochetes/field was observed in the Collagen + NHDFs + HaCaTs (8.7 spirochetes/field) condition compared to the Collagen + HaCaTs (18.3 spirochetes/field) and Collagen Only condition (17.4 spirochetes/field), though these differences were not significant (Fig. 3E). The average number of spirochetes/field were comparable between Collagen + NHDFs + HaCaTs and Collagen + NHDFs (7.5 spirochetes/field) conditions (Fig. 3E). Taken together, dermal fibroblasts limit *B. burgdorferi* growth and colonization of organotypic skin.

It is known that fibroblasts are capable of being invaded by *B. burgdorferi* [32,56–58]. We observed some interaction between NHDFs and *B. burgdorferi,* with as many as 15% of NHDFs associated with bacteria by 48 h post-inoculation (Fig. S4A and B). However, there was minimal evidence of bacterial uptake (Fig. S4A). This confirms previous work with NHDFs that showed little uptake of *B. burgdorferi* in cell culture [57].

### Reactive oxygen species has no effect on RB formation in organotypic skin

One stress *B. burgdorferi* could encounter while invading our organotypic skin model is reactive oxygen species (ROS). Using a 1:1 media mix lowers available sodium pyruvate (NaPyr), which has been shown to increase sensitivity to low concentrations of hydrogen peroxide (H_2_O_2_) [59]. Upon addition of 5 mM NaPyr, we saw no difference in invasion (Fig. 4A and B) and RB formation (Fig. 4A and D). In the presence of our positive control, H_2_O_2_, we saw a statistically significant decrease in *B. burgdorferi* invasion (5.88 Log_2_ fold decrease) (Fig. 4C) while we saw a statistically significant rescue in invasion with 5 mM NaPyr supplementation (0.138 Log_2_ fold change) (Fig. 4A and C). Spirochetes were counted in the media below after 24 h in each condition and no differences in the average number of *B. burgdorferi*/field were observed (Fig. 4E).

**Figure 4.**
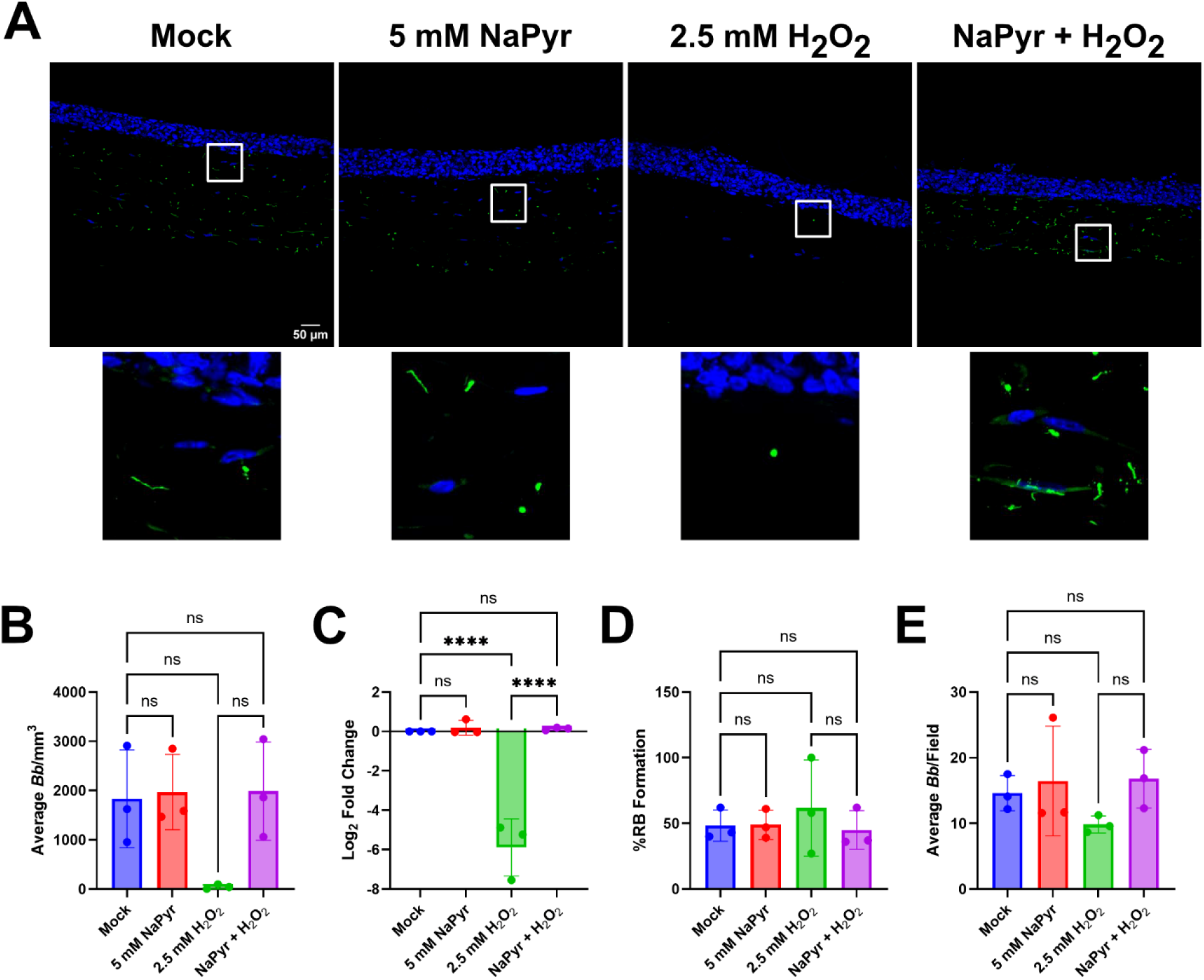
Addition of sodium pyruvate has no effect on total *B. burgdorferi* and %RB formation. (A) Representative images of Mock, 5 mM sodium pyruvate (NaPyr), 2.5 mM hydrogen peroxide (H_2_O_2_), and 5 mM NaPyr + 2.5 mM H_2_O_2_ infection conditions at 24 h. 5 mM NaPyr and 2.5 mM H_2_O_2_ are added at 0 h and serves as a rescue condition. (B) Total *Borrelia* was calculated in each condition at 24 h. (C) Log_2_ fold change of (B) compared to the Mock infection condition. (D) %RB formation was calculated in each condition at 24 h. (E) Spirochetes were counted in the media below in each condition and expressed as Average *Bb*/Field. Experiments were performed in biological triplicate and with technical duplicates. One-way ANOVA with Šídák’s multiple comparisons test was used to determine statistical significance in (B), (C), (D), and (E). Error bars represent standard deviation. Significance is denoted as \*\*\*\**p* < 0.0001.

### Tissue-to-tissue *B. burgdorferi* colonization can be modeled with organotypic skin

Tissue-to-tissue colonization with *B. burgdorferi* has mostly been studied in the context of mice [38–40,42]. Using the SDA system, we were able to colonize a secondary tissue. Briefly, we first colonized organotypic skin in a transwell as described in Figure 2. After 24 h, we washed the skin to remove residual spirochetes and placed the uninfected organotypic skin with the epidermis side down in a new well and covered with 1:1 media. Finally, we place the transwell with the infected skin on top and allow infection to progress for an additional 24 h such that spirochetes could migrate out of the upper skin and into the lower, uninfected skin (Fig. 5A). Using this system, we were able to demonstrate that *B. burgdorferi* can leave the primary infected tissue and infect a secondary tissue below (Fig. 5B).

**Figure 5.**
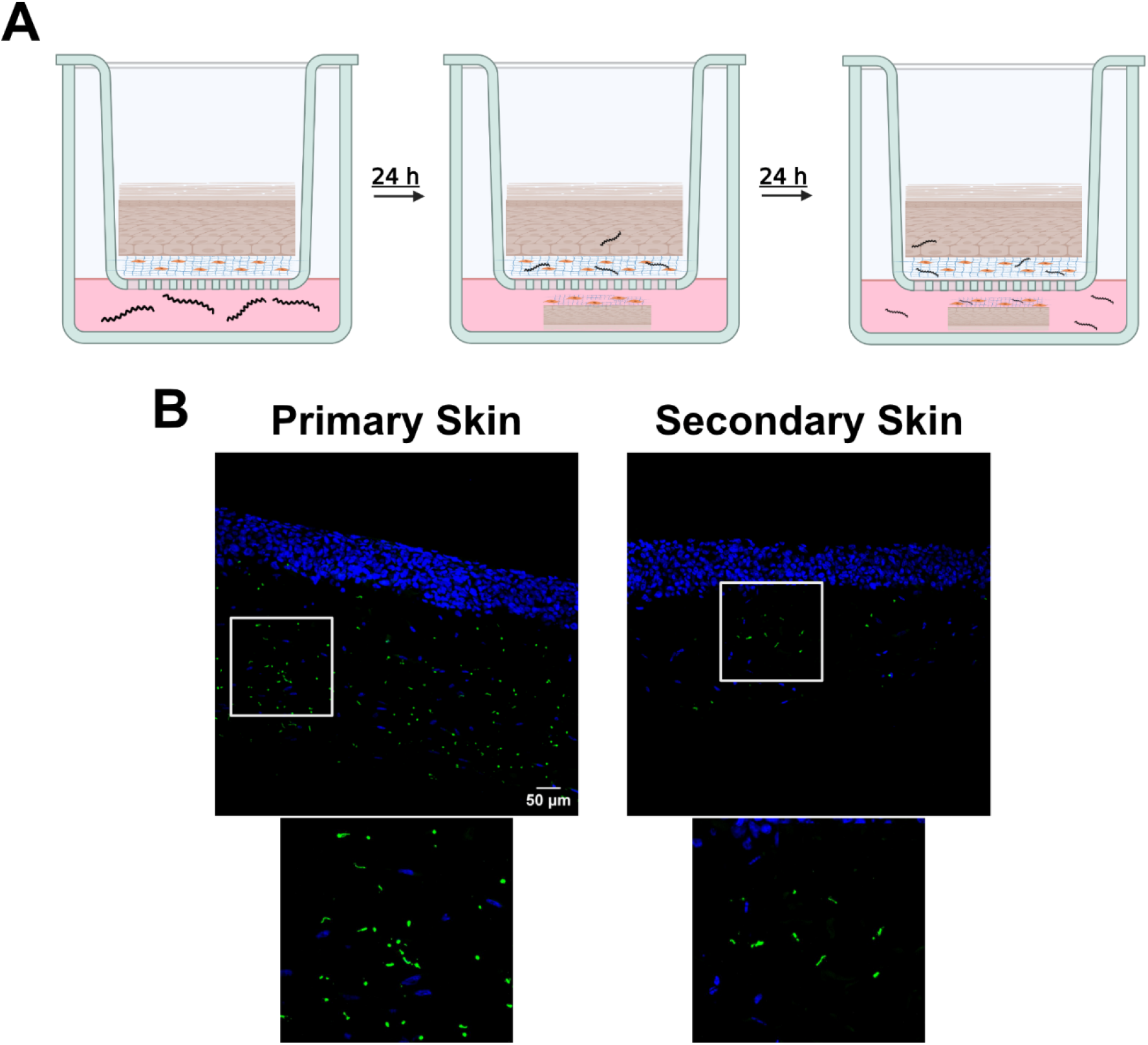
*B. burgdorferi* can colonize secondary organotypic skin. (A) *B. burgdorferi* is placed in 1:1 media for 24 h to allow for colonization of the primary tissue. After 24 h, the primary tissue is moved to a separate well and washed twice with Haley’s Buffer before a secondary tissue is placed beneath and covered with media. An additional 24 h allows for colonization of the secondary tissue. (B) Representative images of the infected primary skin residing in the transwell and the infected secondary skin placed epidermis side down in the media below. DAPI marks nuclei (blue) and anti-*B. burgdorferi* FITC-polyclonal marks *B. burgdorferi* (green).

Overall, we have demonstrated that this model can be colonized in a manner that resembles a bloodstream to skin secondary colonization event. We have observed that dermal fibroblasts play a vital role in the innate immune response to *B. burgdorferi*, resulting in an increase in the RB pleomorphic form. We can also use our organotypic skin model to monitor colonization of a secondary tissue.

## Discussion

Established models to study *B. burgdorferi* infection and dissemination have mostly been limited to murine and *ex vivo* skin models [13,14,17–19,60]. While both useful, they have their limitations. Animal models, murine skin specifically, cannot be directly translated to the events that happen in human skin due to differences in structure, thickness, and cellular content among others [36]. *Ex vivo* human skin models are the gold standard, but tissue availability and patient variability issues can prove difficult for researchers. Herein, we have adapted an organotypic human skin model [29] to study *B. burgdorferi* colonization in a newly developed assay.

A secondary dissemination assay (SDA) was developed to mimic a reinfection event in the skin, which would produce multiple *erythema migrans* beyond the original tick-bite site *in vivo*. This assay revealed the presence of stressed pleomorphic *B. burgdorferi* forms, including round bodies (RBs) (Fig. 2C and F). RBs are a common stress form in response to stimuli, including antibiotics, osmotic shock, serum starvation, and oxidative stress among others [61,62]. This morphological switch has mostly been observed in response to antibiotics [63] and is thought to be linked to Post-Treatment Lyme Disease Syndrome (PTLDS) [62]. This is controversial, as a previous systematic review has found no evidence linking these persistent *B. burgdorferi* forms to chronic LD in humans [64]. There has been some progress made recently, with the development of PTLDS being linked to the presence of persistent forms in the primary infection [65]. Our SDA has shown that RBs are identifiable in organotypic skin, with a peak seen at 48-72 h in a time course (Fig. 2F). At the same time, the majority of *B. burgdorferi* are found in the dermis. (Fig. 2H). Outside of RBs, other pleomorphic forms like “degraded” morphologies and “blebs” have been identified as well (Fig. S2B).

To potentially increase *B. burgdorferi* adhesion and invasion in our model, we added the human basement membrane extract MaxGel, which contains skin relevant components, including glycosaminoglycans, elastin, fibronectin, and laminin. Upon addition of MaxGel, no difference in invasion (Fig. S3A and B) or RB formation was observed (Fig. S3A and C), indicating that collagen in the skin is sufficient for *Borrelia* invasion.

To identify the cause of RB formation, we separated our model into its individual components. In organotypic skin lacking fibroblasts (Collagen Only and Collagen + HaCaTs), more *B. burgdorferi* was observed at 24 h compared to a complete skin condition (Collagen + NHDFs + HaCaTs) (Fig. 3C) while there was a decrease in RB formation compared to complete skin (Fig. 3D). When the fibroblast only (Collagen + NHDFs) condition was compared to the complete skin condition, more *B. burgdorferi* were observed in the complete skin (Fig. 3C) while RB formation was comparable between the two conditions (Fig. 3D). This implicates a dermal fibroblast-mediated mechanism of RB formation.

It is known that dermal fibroblasts are one of the main cells within skin that interact with *B. burgdorferi*. Dermal fibroblasts have been shown to be infected by *B. burgdorferi* [32,57,58,66]. They are also capable of upregulating cytokines and chemokines in response [20,54–56,67,68]. Using our model, we observed interactions between *B. burgdorferi* and NHDFs at 1, 24, and 48 h (Fig. S4D). An increase in interactions was observed up to 24 h, though it was non-significant (Fig. S4E). It is likely that the fibroblast and keratinocyte response to *B. burgdorferi* involves production of cytokines and chemokines in the supernatant below infected organotypic skin, which we plan to identify and evaluate in future studies.

It is possible that antimicrobial peptides (AMPs) produced by fibroblasts, keratinocytes, or both could be contributing to RB formation. Current literature has shown that certain AMPs, such as the human cathelicidin LL-37, rapidly affect *B. burgdorferi* motility [69] but this remains controversial, as other studies note a lack of activity against *B. burgdorferi* [70,71]. Interestingly, fibroblasts were shown to have higher expression of LL-37 in response to *B. burgdorferi*, while keratinocytes had much lower expression of LL-37 and had higher expression of human β-defensin-2 (HBD-2) [72]. When HBD-2 was tested against *B. burgdorferi*, it had lower activity in comparison to LL-37 [69]. Other AMPs have been investigated against *B. burgdorferi* including tick [73] and synthetic AMPs [73,74], with synthetically derived ones showing promise.

Outside of AMPs, oxidative stress could play a role in RB formation in our model. It has already been shown that pyruvate has a protective effect with *B. burgdorferi* and prevents hydrogen peroxide-mediated killing [59]. However, addition of the ROS quencher sodium pyruvate to our model failed to prevent RB formation (Fig. 4D). Alongside oxidative stress, the presence of complement could play a role in *B. burgdorferi* RB formation. Previous studies have shown that fibroblasts highly express complement proteins in *erythema migrans* and control skin biopsies [20], though it is unclear how fibroblasts directly contribute to complement-mediated killing. *B. burgdorferi* strains vary in their complement resistance and employ a variety of methods to avoid complement-mediated killing [75]. These include lipoproteins on their surface as well as hijacking tick saliva complement inhibitors [75].

Using the SDA, we have demonstrated that tissue-to-tissue colonization is possible in our system (Fig. 5B). This sets the groundwork for future studies that includes putting other human tissues in our system, such as heart, joint, and brain tissues which can be colonized by *B. burgdorferi in vivo* [7]. We plan to assess secondary colonization of other organoid and ex vivo tissues in future studies.

A limitation of our current model is the simplicity of it. Future investigations with organotypic skin will include additional cell types that could alter the course of infection, including macrophages, endothelial cells, dendritic cells, and Langerhans cells. The addition of these cell types will allow researchers to tease apart individual resident cell type contributions to *B. burgdorferi* infection either from a primary infection or a secondary dissemination event.

Additionally, the use of HaCaT keratinocytes instead of primary keratinocytes can present issues. It is known that there are deficiencies in the formation of the stratum corneum when using HaCaTs in comparison to primary keratinocytes, though there still is a high degree of epidermal differentiation (Fig. 1) [29]. There can also be differences in the response to cytokine responses between primary and immortalized cell lines, which could present an issue during the immune response to *B. burgdorferi* [76]. Future studies will also include primary keratinocytes in our model to characterize their response to *B. burgdorferi* infection.

Our SDA assay also does not recapitulate early events of *B. burgdorferi* infection following a tick bite, which introduces key saliva factors that are known to alter the host immune response and has been shown to enable efficient *B. burgdorferi* colonization in *ex vivo* and animal inoculation models [17–19,77]. We plan to use this organotypic model to assess which skin-resident cells are altered by tick feeding to enable colonization in future studies.

## Materials and Methods

### Cell line culture conditions and organotypic skin development

Normal Human Dermal Fibroblasts (NHDFs) were cultured at 37°C with 5% CO_2_ using Fibroblast Basal Medium (ATCC, PCS-201-030) with a fibroblast low-serum growth kit (ATCC, PCS-201-041) (FBM). HaCaT keratinocytes were cultured at 37°C with 5% CO_2_ with a previously defined keratinocyte media (Schoop *et al.,* 1999) containing gentamicin. When generating organotypic skin, a collagen mixture containing PBS (Gibco, 10010-023), 1x MEM (Gibco, 11430-030), 4 mg/mL type I collagen (Corning, 354249), 10 ng/mL fibroblast growth factor 2 (FGF2) (R&D Systems, 3718-FB), and 1.25×10^5^/mL NHDFs in FBM were placed in 8 μm transwells and left to solidify for 20-30 minutes at 37°C. After solidifying, 3×10^5^ HaCaTs were placed on the organotypic skin in the 12-well transwell and left submerged for 48 h in HaCaT media to become confluent. Media was thereafter only added to the bottom chamber, allowing 8-13 days of stratification at the air-surface interface. Over the 10-15 days total, a 1:1 solution of FBM and HaCaT media was used and changed every 48 h.

### *Borrelia burgdorferi* growth and infection conditions

*B. burgdorferi* strain B31A3 was grown in complete BSK-H media containing 6% rabbit serum (Millipore Sigma, B8291) at 37°C with 5% CO_2_ to early-mid log phase (2-5×10^7^/mL) as determined by darkfield microscopy. Prior to infection, the skin was washed once with Haley’s Buffer (2.92 g/L NaCl and 4.77 g/L HEPES) above and below. To determine the optimal media for infection, 3×10^6^ spirochetes were inoculated into 4.5 mL of BSK-H, 1:1 BSK-H and FBM, and FBM only in 5 mL culture tubes. Over the course of 10 days, average *Bb*/Field and percent motile spirochetes were calculated. Average *Bb*/Field was calculated by counting the number of spirochetes in each field for a total of 10 fields. Percent motile spirochetes were calculated by counting the number of spirochetes per field that were motile and dividing by the total number of spirochetes in the same field over 10 fields.

### Secondary dissemination assay

For secondary dissemination assays (SDA), 2×10^6^ (4×10^6^ spirochetes/mL) spirochetes were placed underneath the skin in a 1:1 BSK-H:FBM solution and left static at 37°C with 5% CO_2_. For the time course, media was changed after 24 h unless otherwise stated. Prior to fixation at each time point, 10 μL of the media below the skin was removed for counting. 5 randomly imaged fields were counted at each time point or condition. In experiments determining possible causes of round body formation, 5 mM sodium pyruvate and 2.5 mM hydrogen peroxide were added alongside *B. burgdorferi*. After counting, the infected skin was washed once with PBS and fixed using 4% paraformaldehyde for 1 h at room temperature (RT) before being subjected to a sucrose gradient for cross-sectioning. Briefly, the skin was dehydrated stepwise in 15% and 30% sucrose prior to submission to the UNMC Tissue Sciences Facility for OCT embedding, freezing, and generation of ∼30 μm cross-sections on glass slides.

### *Borrelia* invasion by immunofluorescence

Frozen cross-sections were thawed out at room temperature for 1 h prior to blocking and permeabilization (4% BSA, 5% goat serum, and 0.125% Triton X-100 in sterilized PBS + Tween 20 (PBST)) for an additional hour at RT. DAPI (Sigma-Aldrich, 10236276001, 5 μg/mL), FITC-conjugated *B. burgdorferi* Rabbit Polyclonal Antibody (Invitrogen, PA173005, 1:500), and AlexaFluor 647-conjugated wheat germ agglutinin (AF647-WGA) (Invitrogen, W32466, 1 μg/mL) were incubated at RT for 1 h before being washed twice with PBST. Coverslips were mounted using Immu-Mount (Epredia, 9990402) and left overnight to solidify. Images were captured using a 20x objective on a Nikon CSU-W1 spinning disk confocal microscope with 0.9 μm z-slices from 5 independent, random fields per condition in every replicate. Raw images were processed using ImageJ with each condition being its own reference for standardizing image brightness and contrast. Maximum intensity projection images were generated to count *B. burgdorferi* throughout the stack. Images were blinded prior to counting. Total *Borrelia* and round bodies were counted and normalized to the volume of the image. The equation used for normalization is:

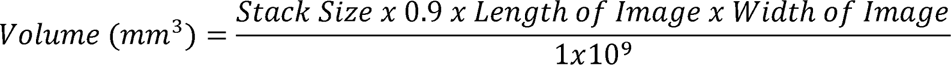

### Quantifying fibroblast*-Borrelia* interactions

Using generated maximum intensity projection images at 1, 24 and 48 h, total fibroblasts were counted based on presence of nuclei marked by DAPI. To confirm location of nuclei in the dermis, AF647-WGA was used. % *B. burgdorferi* positive NHDFs were calculated by counting the total number of fibroblasts that either had *B. burgdorferi* touching or inside the host cell and dividing by the total number of fibroblasts. 5 independent, random fields were analyzed in technical duplicate for a total of 10 fields per biological replicate.

### Tissue-to-tissue dissemination assay

Organotypic skin was infected using the SDA with 2×10^6^ spirochetes (4×10^6^ spirochetes/mL) for 24 h before being moved into a new well where the space below was washed twice with Haley’s Buffer. After washing, a secondary skin sample was placed epidermis side down below the infected skin and covered with 1:1 BSK-H:FBM for an additional 24 h.

Afterwards, the samples were fixed and prepared as previously mentioned for cross-sectioning and staining.

### Statistical analysis

GraphPad Prism version 11.0.2 was used for statistical analysis. A one-way ANOVA with Šídák’s multiple comparisons test was used to determine statistical significance during the SDA time course for selected time points. Two-way ANOVA with Tukey’s multiple comparisons test was used to compare the number of *B. burgdorferi* and RB formation in the dermis and epidermis across the time course. One-way ANOVA with Dunnett’s multiple comparisons test was used to determine statistical significance between different organotypic skin compositions while a Welch’s *t* test was used to determine significance between the NHDF only and NHDFs + HaCaTs groups. One-way ANOVA with Šídák’s multiple comparisons test was used to determine significance upon the addition of MaxGel, sodium pyruvate, and hydrogen peroxide in different organotypic skin conditions. Graphics were generated using BioRender and figures compiled using Affinity Designer 2.

## Contributions and Acknowledgements

We would like to thank the laboratories of Dr. Sujata Chaudhari and Dr. Rey Carabeo for their invaluable feedback regarding this project. We would especially like to thank Dr. Rey Carabeo for sharing his laboratories’ space and equipment that enabled us to begin this project.

JJK performed the experiments in this manuscript, while JJK and AJB planned the experiments, wrote and edited the manuscript. The funding supporting this work was awarded to AJB through the following mechanisms: NIH/NIAID R21AI185574-01A1, the Mary G. and George W. White Fund for Medical Research Award, and the Great Plains IDEA CTr Early Career Investigator Award (NIGMS,U54-GM115458).

---

**Supplemental Figure 1.**
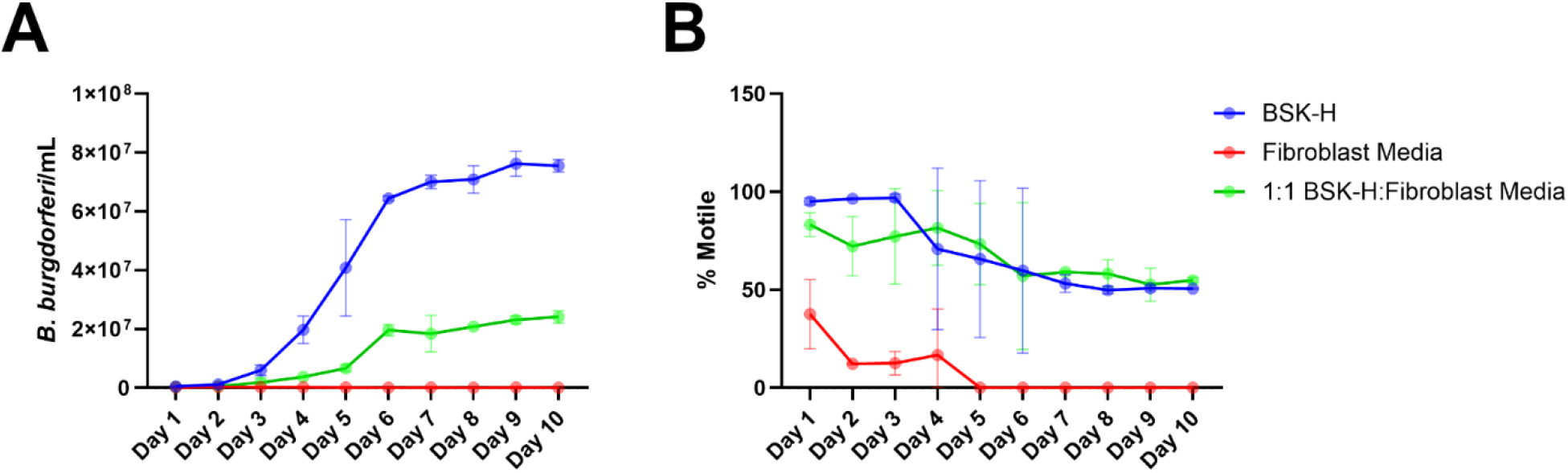
Combination BSK-H:Fibroblast media supports *B. burgdorferi* growth and motility. (A) Across 10 days, 10 fields were counted per coverslip in each media condition. (B) Motility was assessed by counting the number of motile spirochetes in each field and dividing by the total number generated from 10 fields in (A). Two biological replicates are shown in (A) and (B).

**Supplemental Figure 2.**
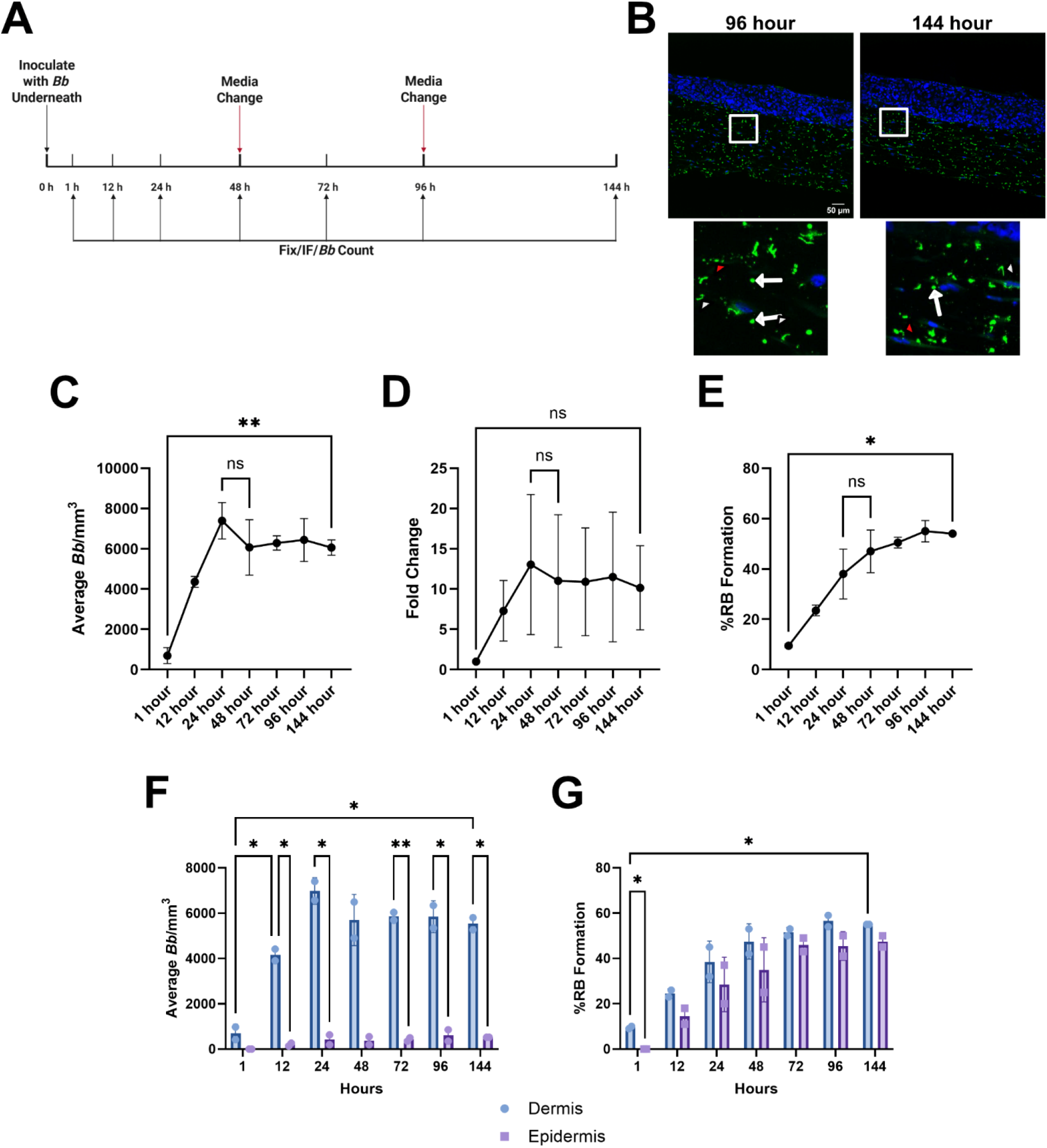
*B. burgdorferi* is present in organotypic skin out to 144 h. (A) Timeline of infection. At each indicated time point, infected skin was fixed, stained for *B. burgdorferi*, and counted. Media was changed at 48 and 96 h to sustain skin viability. (B) Representative images of 96 and 144 h time points. White arrows point to RBs, white arrowheads point to blebs, and red arrowheads point to degraded forms. (C) Total *Borrelia* were calculated at each time point. (D) Fold change of total *Borrelia* compared to 1 h in (C). (E) %RB formation was calculated at each time point. (F) Total *Borrelia* was calculated for the dermis and epidermis to determine localization. (G) %RB formation was calculated at each time point to determine localization. For all graphs, only statistically significant comparisons are shown. Experiments were performed in biological duplicate and with technical duplicates. Each dot represents the average of a biological replicate. One-way ANOVA with Šídák’s multiple comparisons test was used to determine statistical significance in (C), (D), and (E). Two-way ANOVA with Tukey’s multiple comparisons test was used to compare each time point and location for (F) and (G). Error bars represent standard deviation. Significance is denoted as \**p* < 0.05 and \*\**p* < 0.01.

**Supplemental Figure 3.**
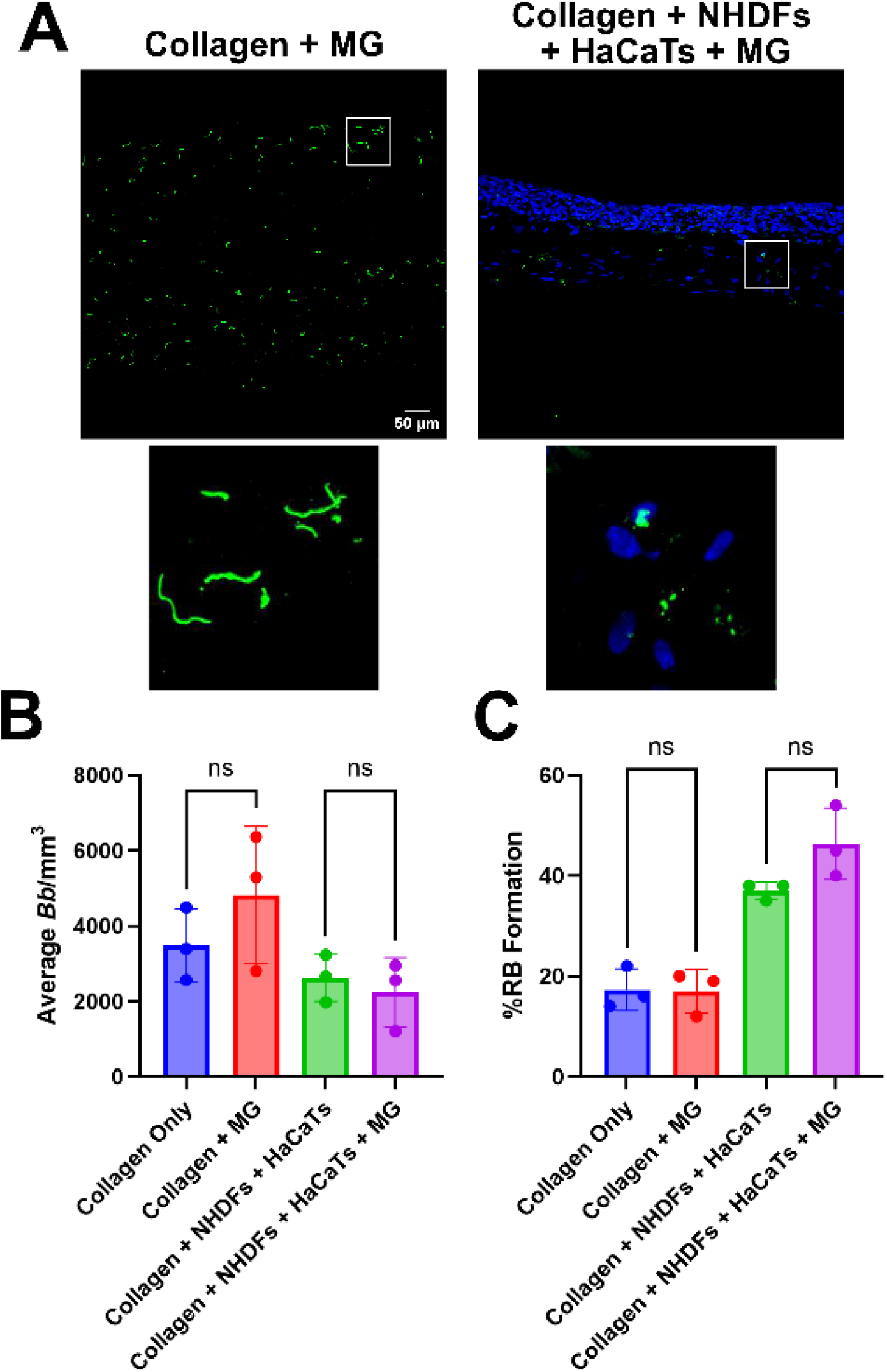
Addition of skin-associated ECM components have no effect on colonization. (A) MaxGel (1:1000) was added to two conditions: Collagen Only and complete organotypic skin (Collagen + NHDFs + HaCaTs). Representative images are shown for both conditions. (B) Total *Borrelia* was calculated in each condition at 24 h across 5 fields and normalized to the total captured area. (C) %RB formation was calculated in each condition at 24 h. Experiments were performed in biological triplicate and with technical duplicates. One-way ANOVA with Šídák’s multiple comparisons test was used to determine statistical significance in (B) and (C). Error bars represent standard deviation.

**Supplemental Figure 4.**
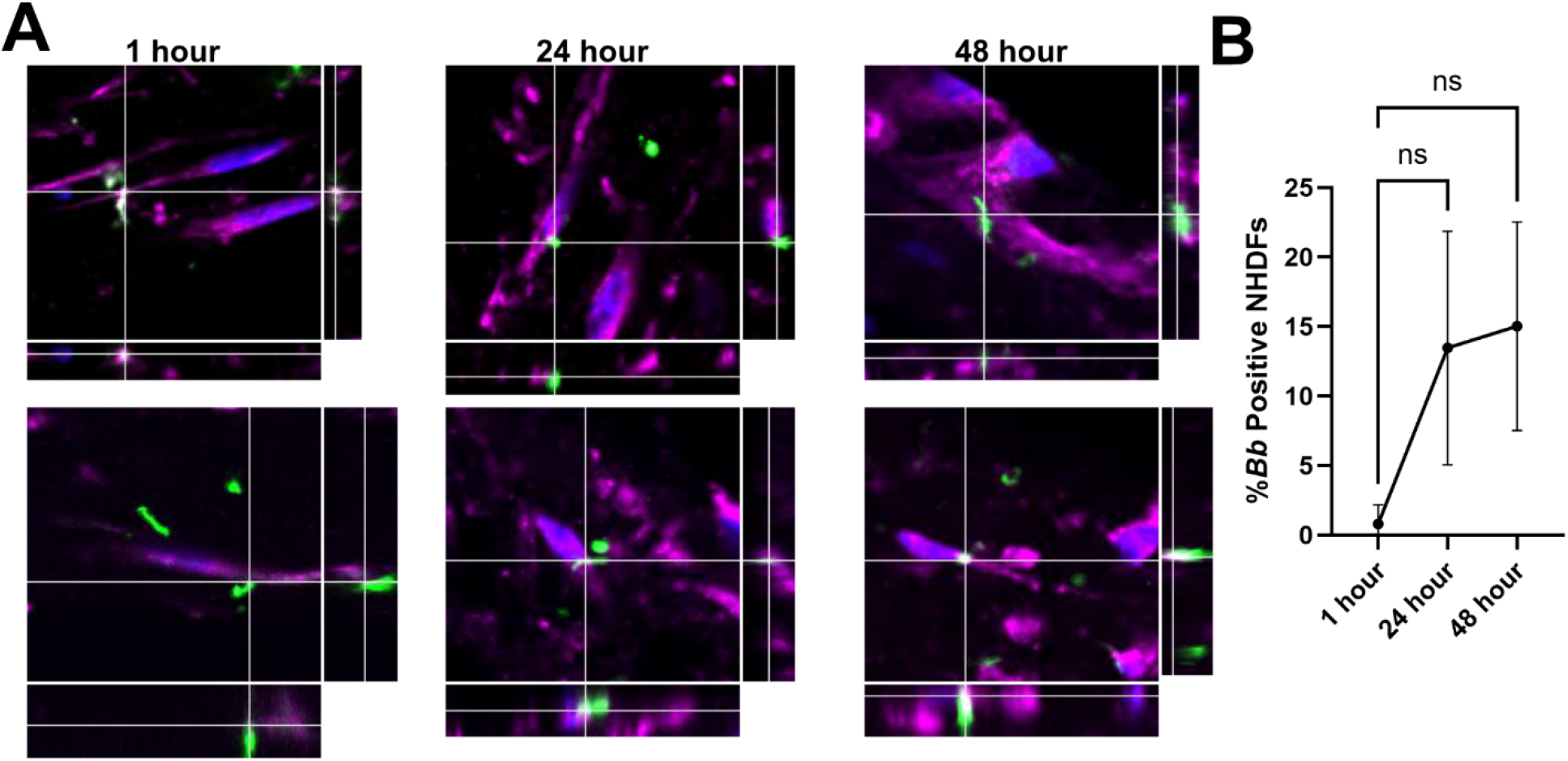
NHDFs interact with *B. burgdorferi* in organotypic skin. (A) Two representative images of NHDFs interacting with *B. burgdorferi* at 1, 24, and 48 h in a single z-slice. DAPI marks nuclei (blue), anti-*B. burgdorferi* FITC-polyclonal marks *B. burgdorferi* (green), and WGA marks fibroblasts (magenta). (B) % *B. burgdorferi* positive NHDFs were calculated at each time point. A positive NHDF was defined as either *B. burgdorferi* touching or inside the fibroblast. One-way ANOVA with Dunnett’s multiple comparisons test was used to determine statistical significance in (B). Error bars represent standard deviation.

**Supplemental Figure 5.**
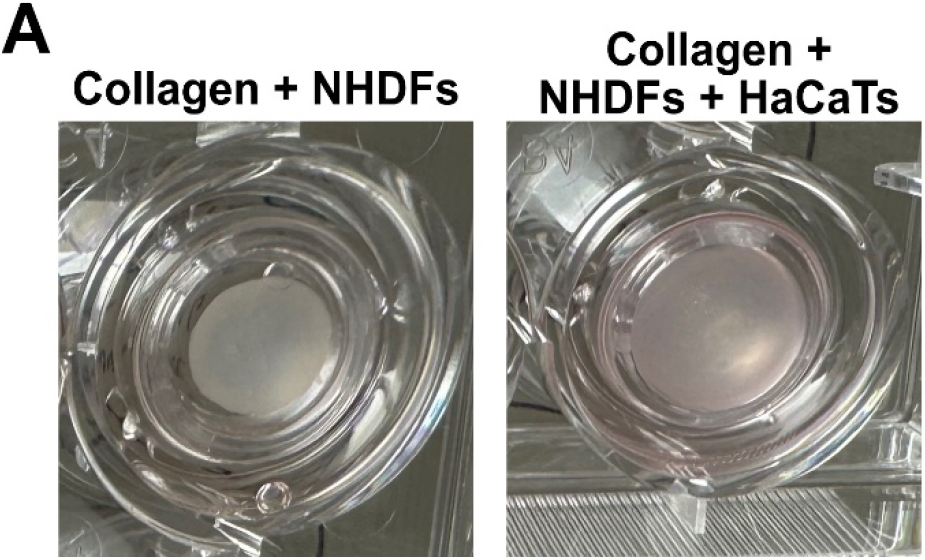
Fibroblast only organotypic skin is shrunken. (A) Representative images of Collagen + NHDFs and Collagen + NHDFs + HaCaTs skin conditions, indicating fibroblast-mediated shrinking of the organotypic skin.

